# Cardiac fibroblast GSK-3α mediates adverse myocardial fibrosis via IL-11 and ERK pathway

**DOI:** 10.1101/2021.02.02.429435

**Authors:** Prachi Umbarkar, Sultan Tousif, Anand P. Singh, Joshua C. Anderson, Qinkun Zhang, Hind Lal

## Abstract

**Background:** Heart failure is the leading cause of mortality, morbidity, and healthcare expenditures worldwide. Numerous studies have implicated Glycogen Synthase Kinase-3 (GSK-3) as a promising therapeutic target for cardiovascular diseases. GSK-3 isoforms appear to play overlapping, unique, and even opposing functions in the heart. Recently our group has identified cardiac fibroblast (CF) GSK-3β as a negative regulator of fibrotic remodeling in the ischemic heart. However, the role of CF-GSK-3α in myocardial fibrosis is unknown.

**Methods and Results:** Herein, we employed two entirely novel conditional fibroblast-specific and tamoxifen-inducible mouse models to define the role of CF-GSK-3α in fibroblast activation and myocardial fibrosis. Specifically, GSK-3α was deleted from cardiac fibroblasts or myofibroblasts with tamoxifen-inducible Tcf21- or periostin-promoter-driven Cre recombinase. At 2 months of age, WT and KO mice were subjected to cardiac injury, and heart functions were monitored by serial echocardiography. Histological analysis and morphometric studies were performed at 8 weeks post-injury. In both settings, GSK-3α deletion restricted fibrotic remodeling and improved cardiac function. To investigate underlying mechanisms, we examined the effect of GSK-3α deletion on myofibroblast transformation and pro-fibrotic TGFβ1-SMAD3 signaling *in vitro*. A significant reduction in cell migration, collagen gel contraction, and α-SMA expression in TGFβ1 treated GSK-3α KO MEFs confirmed that GSK-3α is required for myofibroblast transformation. Surprisingly, GSK-3α deletion did not affect SMAD3 activation, indicating the pro-fibrotic role of GSK-3α is SMAD3 independent. To further delineate the underlying mechanism, total proteins were isolated from CFs of WT and KO animals at 4 weeks post-injury, and kinome profiling was performed by utilizing PamStation^®^12 high throughput microarray platform. The kinome analysis identified the downregulation of RAF family kinase activity in GSK3α-KO-CFs. Moreover, mapping of significantly altered kinases against literature annotated interactions generated ERK-centric networks. Importantly, flow cytometric analysis of CFs confirmed a significant decrease in pERK levels in KO mice. Additionally, our *in vitro* studies demonstrated that GSK-3α deletion prevented TGFβ1 induced ERK activation thereby validating our findings from kinome analysis. Interestingly, IL-11, a fibroblast specific downstream effector of TGFβ1, was very low in GSK-3α KO MEFs as compared to WT and ERK inhibition further reduced IL-11 expression in them. All these results indicate that GSK-3α mediates pro-fibrotic response in the injured heart through IL-11 and ERK pathway.

**Conclusion:** CF-GSK-3α plays a causal role in myocardial fibrosis that could be therapeutically targeted for future clinical applications.

## Introduction

HF is a leading cause of death and a major socioeconomic burden for societies across the world. Most heart diseases are associated with fibrosis. ^1–3^ However, there is no specific therapy to combat myocardial fibrosis. The absence of direct evidence at clinical and pre-clinical levels strengthens the notion that fibrosis is not a cause of the disease; rather, it is a consequence of the pathological events occurring in the failing heart. Cardiac fibroblast (CF), one of the major cell types in the heart ^4^, is a key contributor to the development of fibrosis.^5^ Until recently, the role of CF in cardiac pathophysiology remained auxiliary to cardiomyocyte. Primarily because animal models that permit genetic manipulation of target gene specifically in fibroblasts were not generated. Thus, most of the literature regarding fibroblast biology has largely relied on experimental outcomes emerging from *in vitro* culture systems and mouse models in which the target gene is deleted in cardiomyocytes or globally. To overcome this limitation, the scientific community has developed fibroblast-specific genetic mouse models.^6^ Among the recently generated FB-specific genetic models, Tcf21^MCM^ and Postn^MCM^ mice lines are well characterized for the effective manipulation of a target gene in the tissue-resident fibroblasts and myofibroblasts, respectively.^7–13^

Myocardial fibrosis is characterized by the disproportionate accumulation of extracellular matrix (ECM) components. In an injured heart, CFs undergo a phenotypic switch from quiescent to an activated state. These activated CFs or myofibroblasts are considered as principal producers of ECM under pathological conditions.^5^ Among various pro-fibrotic factors, TGFβ1 is the best characterized for its role in CF activation and has been implicated with fibrosis in various cardiac diseases. In canonical TGFβ signaling, binding of TGFβ to its receptors leads to activation of receptor-associated SMADs. These activated SMADs form a heteromeric complex with co-SMAD i.e. SMAD4 which then translocates into the nucleus and regulates transcription of genes. In addition to the canonical signaling, TGFβ can modulate signaling of mitogen-activated protein kinases (MAPKs), phosphoinositide-3-kinase (PI3K), Ras Homolog Family Member A (RhoA), protein phosphatase 2A (PP2A), nuclear factor-κB (NF-kB), and TGF-β-activated kinase 1 (TAK1). Our lab has demonstrated the CF-specific deletion of GSK-3β causes hyperactivation of SMAD-3, which results in excessive fibrosis and cardiac dysfunction post-MI.^15^ Also, fibroblast-specific genetic manipulation revealed that MAPK p38α acts as a nodal point where mechanical and paracrine signals converge to initiate a fibrotic response in the mouse heart.^10^ A recent study has shown that TGF β1 increases IL-11 expression in fibroblast which in turn regulates translation of fibrogenic proteins via non-canonical, ERK-dependent pathway.^16^ All these studies indicate that multiple distinct pathways cooperate with classical TGF β signaling in fibroblast to mediate fibrotic response in the diseased heart.

The Glycogen synthase kinase-3 (GSK-3) is a family of Serine/Threonine kinases. It has two highly conserved isoforms, GSK-3α and GSK-3β. The roles of GSK-3s in cardiomyocyte biology and cardiac disease are extensively studied.^15, 17–21^. Our lab has discovered that CF-GSK-3β acts as a negative regulator of myocardial fibrosis in the ischemic heart.^15^ However, the role of CF-GSK-3α in the pathogenesis of heart failure is completely unknown. In the present study, we employed Tcf21^MCM^ and Postn^MCM^ models to delete GSK-3α specifically from resident cardiac fibroblasts and myofibroblasts. We report that CF-specific deletion of GSK-3α prevents pressure overload-induced adverse fibrotic remodeling and cardiac dysfunction. Mechanistically, we demonstrate that GSK-3α exerts its pro-fibrotic effect via the IL-11 and RAF-ERK pathway and independent of classical canonical TGF-β1/SMAD3 signaling.

## Material and Methods

### Mice

To achieve conditional deletion of GSK-3α specifically in resident fibroblast, GSK-3α ^fl/fl^ mice ^22, 23^ were crossed with mice carrying the tamoxifen (TAM) inducible TCF-21 promoter-driven Mer-Cre-Mer (MCM) transgene (a gift from Michelle D. Tallquist, University of Hawaii, USA). To generate myofibroblast specific GSK-3α KO, GSK-3α ^fl/fl^ mice were crossed with Periostin-MCM mice^11^ (Jackson laboratories, stock #029645). The GSK-3α ^fl/fl^/Cre^+/-^/TAM mice were the conditional knockouts (KO), whereas littermates GSK-3α ^fl/fl^/Cre^-/-^/TAM represented as controls (WT). The Institutional Animal Care and Use Committee of the University of Alabama at Birmingham approved all animal procedures and treatments used in this study (protocol # IACUC-21701).

### Transverse Aortic Constriction (TAC) surgery in mice

TAC was performed as previously described.^22, 24^ Briefly, mice were sedated with isoflurane (induction, 3%; maintenance, 1.5%) and anesthetized to a surgical plane with intraperitoneal ketamine (50 mg/kg) and xylazine (2.5 mg/kg). Anesthetized mice were intubated, and a midline cervical incision was made to expose the trachea and carotid arteries. A blunt 20-gauge needle was inserted into the trachea and connected to a volume-cycled rodent ventilator on supplemental oxygen at a rate of 1 l/min, with a respiratory rate of 140 breaths/min. Aortic constriction was performed by tying a 7-0 nylon suture ligature against a 27-gauge needle. The needle was then promptly removed to yield a constriction of approximately 0.4 mm in diameter. To confirm the efficiency of TAC surgery, the pressure gradient across the aortic constriction was measured by Doppler echocardiography.

### Echocardiography

Echocardiography was performed as described previously (PMID: 30847418, 26976650). In brief, transthoracic M-mode echocardiography was performed with a 12-MHz probe (VisualSonics) on mice anesthetized by inhalation of isoflurane (1-1.5%). LV end systolic interior dimension (LVID;s), LV end diastolic interior dimension (LVID;d), ejection fraction (EF), and fractional shortening (FS) values were obtained by analyzing data using the Vevo 3100 program.

### Histology

Heart tissues were fixed in 10% neutral buffered formalin, embedded in paraffin, and sectioned at 5 μm thickness. Sections were stained with Hematoxylin-Eosin (HT1079, Sigma-Aldrich) or Masson Trichrome (HT15, Sigma-Aldrich) as per the manufacturer’s instructions. The images of the LV region were captured using a Nikon Eclipse E200 microscope with NIS element software version 5.20.02. The quantification of LV fibrosis and cross-sectional area of cardiomyocytes (CSA) were determined with ImageJ version 1.52a software (NIH). At least 200-300 cardiomyocytes per heart (n=5 hearts per group) were taken for CSA measurement.

### Western blot

LV tissues were dissected from mouse heart and homogenized with cell lysis buffer (Cell signaling Technology) having 50 mM Tris-HCl (pH7.4), 150 mM NaCl, 1mM EDTA, 0.25% sodium deoxycholate, 1% NP-40, and freshly supplemented Protease Inhibitor Cocktail (Sigma-Aldrich) and Phosphatase Inhibitor Cocktail (Sigma-Aldrich) present. Nuclear and cytoplasmic fractions were prepared using NE-PER Reagent from Pierce (catalog #78833) as per the manufacturer’s instructions. Protein concentration was determined with Bradford assay (Bio-Rad protein assay kit). An equal amount of proteins was denatured in SDS–PAGE sample buffer, resolved by SDS–PAGE, and transferred to Immobilon-P PVDF membrane (EMD Millipore). The membranes were blocked in Odyssey blocking buffer (LI-COR Biosciences) for 1h at RT. Primary antibody incubations were performed at different dilutions as described in the antibody list (Supplemental Table 1). All incubations for primary antibodies were done overnight at 4°C and followed by secondary antibody (IRDye 680RD or IRDye 800CW from LI-COR Biosciences) incubation at 1:3000 dilutions for 1h at RT. Proteins were visualized with the Odyssey Infrared Imaging System (LI-COR Biosciences). Band intensity was quantified by Image Studio version 5.2 software.

### RNA extraction and quantitative PCR analysis

Total RNA was extracted from heart tissue using the RNeasy Mini Kit (74104, Qiagen) according to the manufacturer’s protocol. cDNA was synthesized using the iScript cDNA synthesis kit (170-8891, Bio-Rad) following the manufacturer’s instructions. Gene expression was analyzed by quantitative PCR (qPCR) using the TaqMan Gene Expression Master Mix (4369016, Applied Biosystems) and TaqMan gene expression assays (Applied Biosystems) on a Quant studio 3 (Applied Biosystems) Real-Time PCR Detection machine. Details of TaqMan gene expression assays used in this study is provided as Supplemental Table 2. Relative gene expression was determined by using the comparative CT method (2^-ΔΔCT^) and was represented as fold change. Briefly, the first ΔCT is the difference in threshold cycle between the target and reference genes: ΔCT= CT (a target gene X) - CT (18S rRNA) while ΔΔCT is the difference in ΔCT as described in the above formula between the CTL and KO group, which is = ΔCT (KO target gene X) - ΔCT (CTL target gene X). Fold change is calculated using the 2^-ΔΔCT^ equation.

### Isolation and culture of fibroblasts

#### Adult mouse cardiac fibroblasts

Primary adult mouse cardiac fibroblasts were isolated according to a previous protocol.^25^ Briefly, hearts were excised, rinsed in cold Krebs-Henseleit (Sigma) buffer supplemented with 2.9mM CaCl2 and 24mM NaHCO3. The tissue was minced and transferred to Enzyme cocktail [0.25 mg/mL Liberase TH (Roche), 20 U/mL DNase I (Sigma Aldrich), 10 mmol/L HEPES (Invitrogen), in HBSS] and serial digestion was performed at 37 °C for 30 min. After each digestion, digests were passed through a 40 μm filter and collected in a tube containing DMEM-F12 with 10% FBS. At the end of the digestion protocol, cells were pelleted by centrifugation at 1000 rpm for 10 min. To remove RBCs, the cell pellet was resuspended in 1 ml of RBC/ACK lysis buffer (Gibco) and incubated for 1 min. Following incubation, cells were washed with the KHB buffer and centrifuged at 1000 rpm for 10 min. Cells were resuspended in DMEM-F12 with 10% FBS and plated into a 100mm dish for 2h. Unattached and dead cells were washed with DPBS and fresh media was added to the attached fibroblast for culture.

#### Neonatal Rat Ventricular Fibroblasts - NRVF

Primary cultures of neonatal rat ventricular fibroblasts (NRVFs) were prepared from 1-2-day-old Sprague-Dawley rats as described previously.^15^

#### Mouse embryonic fibroblasts - MEFs

The creation of WT and GSK-3α KO MEFs used in this study have been described previously.^26^ MEFs were grown in Dulbecco’s Modified Eagle’s medium (DMEM) supplemented with 10% fetal bovine serum (GIBCO) and 1% penicillin-streptomycin (GIBCO).

### Adenovirus infection

For adenoviral infection of cardiac fibroblasts, we employed replication-defective adenoviruses. Adenovirus expressing LacZ was used to control for nonspecific effects of adenoviral infection. Levels of expressed proteins were determined by Western blot analysis. For each adenovirus employed, cardiac fibroblasts were treated with viruses at 25, 50, 100, 150, and 200 MOI to maximize protein expression, but to limit viral toxicity. At 24 h after plating, cardiac fibroblasts were infected with adenovirus diluted in DMEM medium. After 24h of adenoviral infection, the medium was replaced with virus-free serum free medium, and cells were cultured for an additional 12 h before experiments. Viral MOI was determined by dilution assay in HEK-293 cells grown in 6-well clusters.

### Cell culture treatments and functional assays

Cells were serum-starved overnight and treated with the following as per experimental needs - ERK inhibitor U0126 (sigma, 5μM), PD98059 (sigma, 10μM), TGFβ1 (Sigma, 10ng/ml) for the indicated period of time. In collagen gel contraction assays, collagen gels were populated with MEFs. Free-floating gels were maintained in culture medium and the gel area was measured at 48h of treatment. In cell migration assays, MEFs were seeded in culture dishes and allowed to grow till confluency. The wound was created by scratching the monolayer of the cells. Cells were then treated with TGF-β1. The percentage of wound closure was monitored and quantified over the indicated time points.

### Kinome analysis

For Kinome studies adult mouse cardiac fibroblasts were isolated from WT and KO animals at 4 weeks post-TAC as described in the previous section. To enrich the FB population, endothelial (CD31+) and myeloid (CD45+) cells were sorted using the Magnetic Cell Isolation and Cell Separation kit (Miltenyi Biotec) with recommended dilutions of antibodies against CD31 (Miltenyi Biotec 130-097-418) and CD45 (Miltenyi Biotec 130-052-301) as per the manufacturer’s instructions. Cells were lysed in an M-Per lysis buffer containing Halt’s protease and phosphatase inhibitors (Pierce). For each sample, 2μg of protein was loaded onto Serine/Threonine Kinase PamChip along with standard kinase buffer (Supplied by PamGene), 100 μM ATP, and FITC-labeled anti-phospho-serine/threonine antibodies. Kinomic profiling was carried out using the PamStation®12 Platform Technology (PamGene, International, Den Bosch, The Netherlands) within the UAB Kinome Core (www.kinomecore.com) as described previously.^27^

### Flow cytometry

CFs were isolated as described in the previous section. For analysis, cells were incubated with Fc blocker followed by fluorochrome-conjugated antibody staining for the following: MEFSK4, CD45, CD31, pERK, IL-11 for 30 min. Details of antibodies used in this study is provided as Supplemental Table 3. Dead cells were excluded using 7AAD or Ghost dyeTM violet 510. CFs were identified as CD45^-^CD31^-^ MEFSK4^+^ cells. Fluorescence intensity of fluorochrome-labeled cells was assessed by flow cytometry (BD LSR-II) and FACS Diva software, and final data analysis was performed by Flow Jo (Tree Star, USA) software.

### Statistical analysis

Analyses were performed using GraphPad Prism (version 9.0.0). Differences between the 2 data groups were evaluated for significance by the Mann-Whitney test. For comparisons of more than 2 groups, ANOVA followed by Tukey’s multiple comparison test was applied. All data are expressed as mean ± SEM. For all tests, a p-value of < 0.05 was considered statistically significant.

## Results

### Deletion of GSK-3α in residential FBs prevents adverse cardiac remodeling & maintains cardiac function

To evaluate the role of cardiac resident fibroblast GSK-3α in injury-induced cardiac remodeling, we generated a mouse model in which GSK-3α was conditionally deleted in resident fibroblast using tamoxifen (TAM)-inducible Tcf21 promoter-driven MerCreMer transgene (Tcf21-MCM). We crossed Tcf21-MCM mice with GSK-3α ^fl/fl^ mice to obtain GSK-3α^fl/fl^ Tcf21^MCM^ mice. GSK-3α^fl/fl^/MCM^+/-^/TAM mice were conditional knockout (KO), whereas littermates GSK-3α ^fl/fl^/MCM^-/-^/TAM represented wild type controls (WT). The tamoxifen (TAM) chow diet protocol was employed to induce the FB-specific expression of Cre recombinase, as previously reported.^8^ To confirm the CF-specific deletion of GSK-3α, fibroblasts were isolated from the heart of experimental animals and proteins were extracted. Western blot analysis confirmed that TAM treatment led to a significant reduction of GSK-3α protein in the CFs from KO mice compared with littermate controls **(Figure 1A–1B)**.

**Figure 1:**
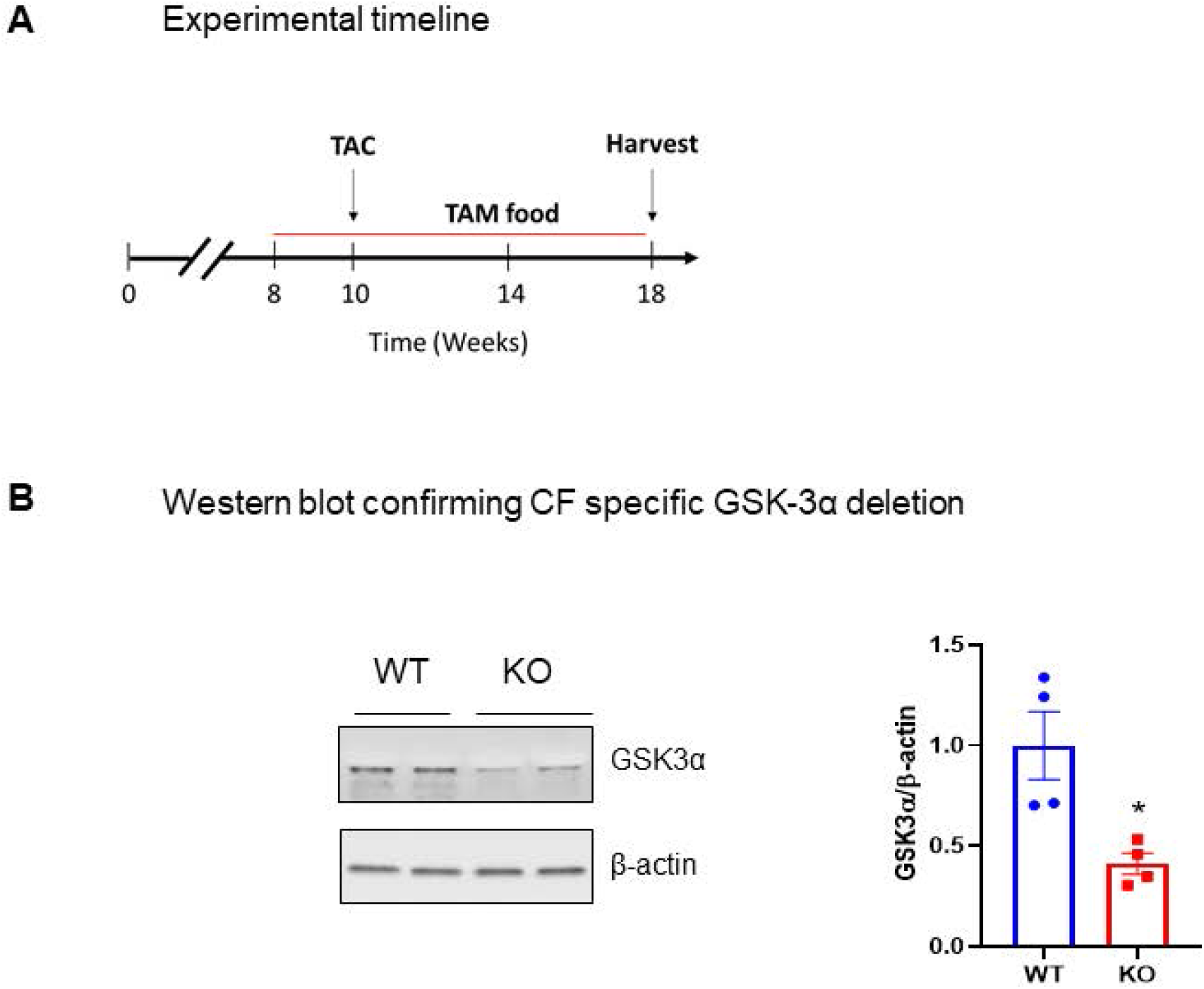
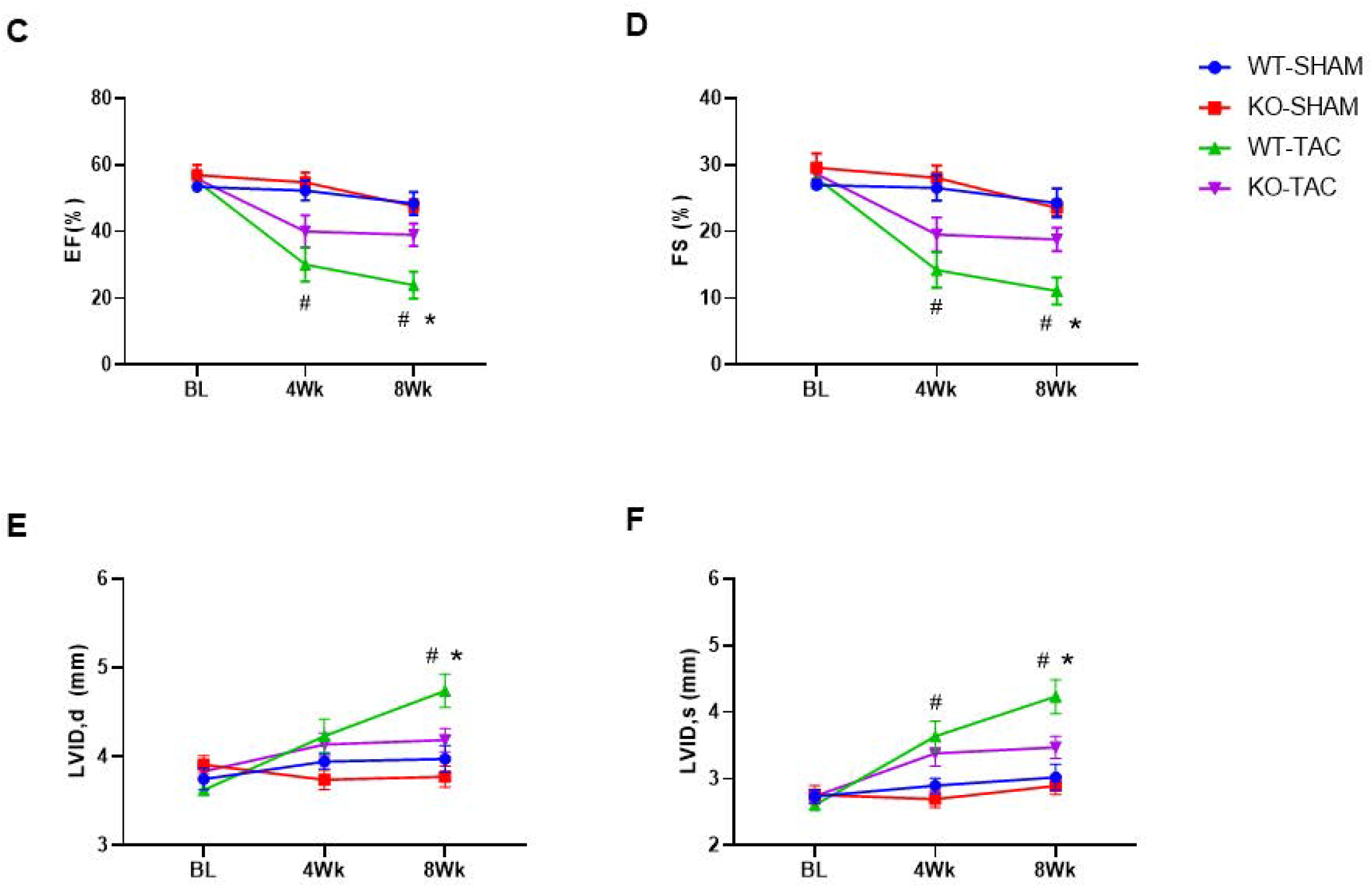
Deletion of CF-GSK-3α from resident cardiac fibroblasts prevents pressure overload-induced cardiac dysfunction. **(A)** Experimental design. Two months old mice were fed the tamoxifen (TAM) chow diet. After 2 weeks of TAM treatment, mice were subjected to TAC surgery and are maintained on the TAM diet till the end of the study. **(B)** Western blot analysis of GSK-3α protein levels in cardiac fibroblasts at 4 weeks after TAM treatment. Data were analyzed using the nonparametric Mann-Whitney test and represented as mean ± SEM. N=4 per group. \**P*<0.05. Evaluation of cardiac function by m-mode echocardiography; **(C)** Ejection fraction (EF), **(D)** Fractional shortening (FS), **(E)** LV end diastolic interior dimension (LVID;d) and **(F)** LV end-systolic interior dimension (LVID;s). Data were analyzed using one-way ANOVA followed by Tukey’s *post hoc* analysis and represented as mean ± SEM. N=5-8 per group. ^#^*P*<0.05 for WT-SHAM vs WT-TAC, \**P*<0.05 for WT-TAC vs.KO-TAC.

After 2 weeks of TAM treatment, mice were subjected to trans-aortic constriction (TAC) surgery. To assess the cardiac function, we performed serial M-mode echocardiography. In the WT-TAC group, LVEF and LVFS started to decline from 4 weeks post-surgery, indicating systolic dysfunction **(Figure 1C–1D)**. These changes were associated with the development of structural remodeling as reflected by a significant increase in LVIDs, in the WT-TAC group **(Figure 1E–1F)**. Interestingly, all these parameters were remarkably normalized in GSK-3α KO. To verify the effects of CF specific GSK-3α deletion on adverse cardiac remodeling, we did morphometrics and histological studies at 8 weeks post-TAC. We measured HW/TL ratio and cardiomyocyte cross-sectional area (CSA). Both these parameters were elevated in the WT-TAC group revealing hypertrophic remodeling **(Figure 2A–2B)**. For assessment of cardiac fibrosis, we did Masson’s Trichrome staining and found excessive collagen deposition in WT-TAC hearts **(Figure 2C–2D).** All these characteristics of pathological cardiac remodeling were alleviated in GSK-3α KO mice. Additionally, we compared expression levels of key genes related to cardiac hypertrophy (ANP and BNP) and fibrosis (COL1A1 and COL3A1). These molecular markers were significantly augmented in the WT-TAC group and were normalized in GSK-3α KO mice **(Figure 2E–2H).** Taken together, these findings confirmed that deletion of GSK-3α from resident FBs before injury insult resulted in improved cardiac function and prevented adverse cardiac remodeling in TAC mice.

**Figure 2:**
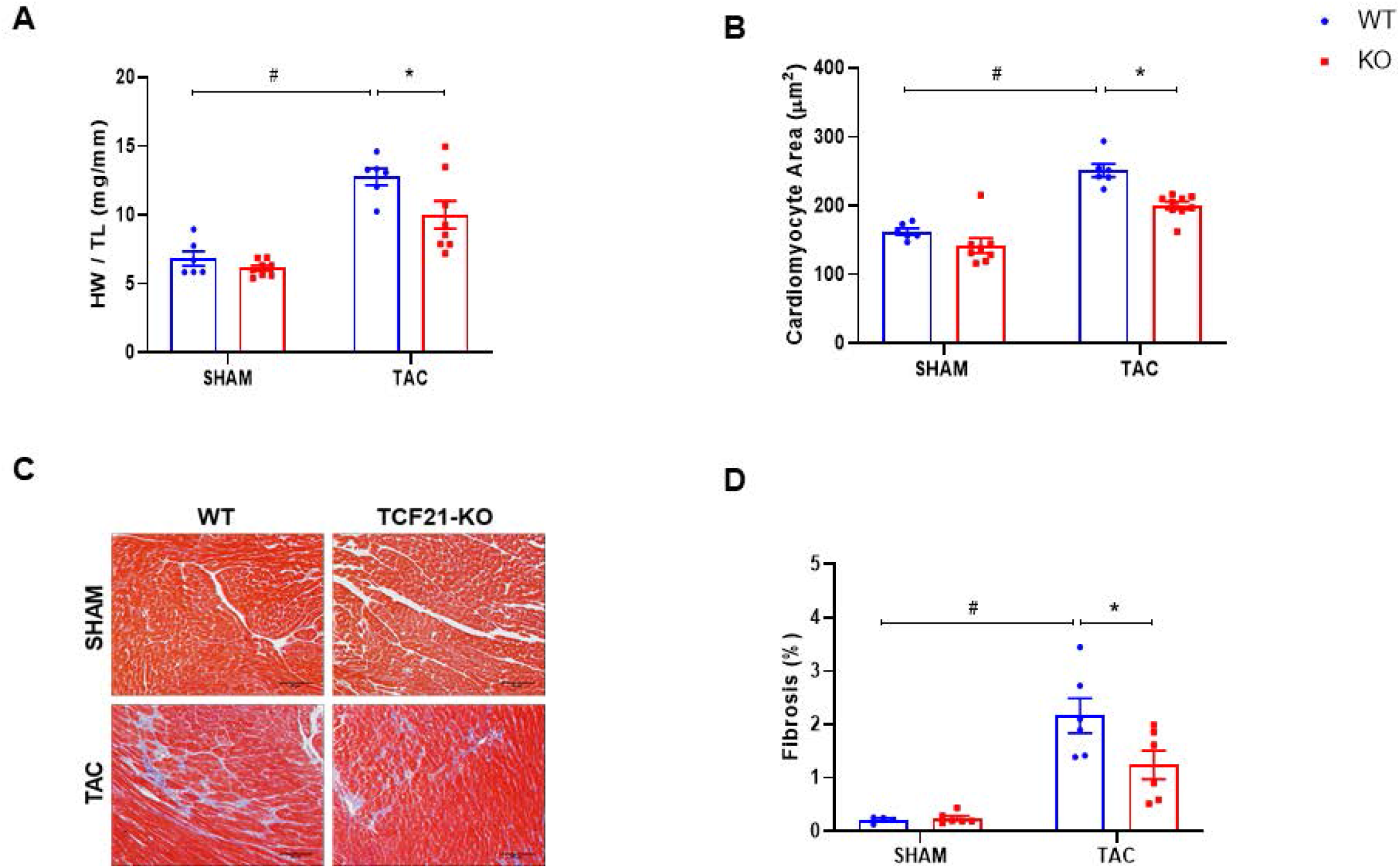
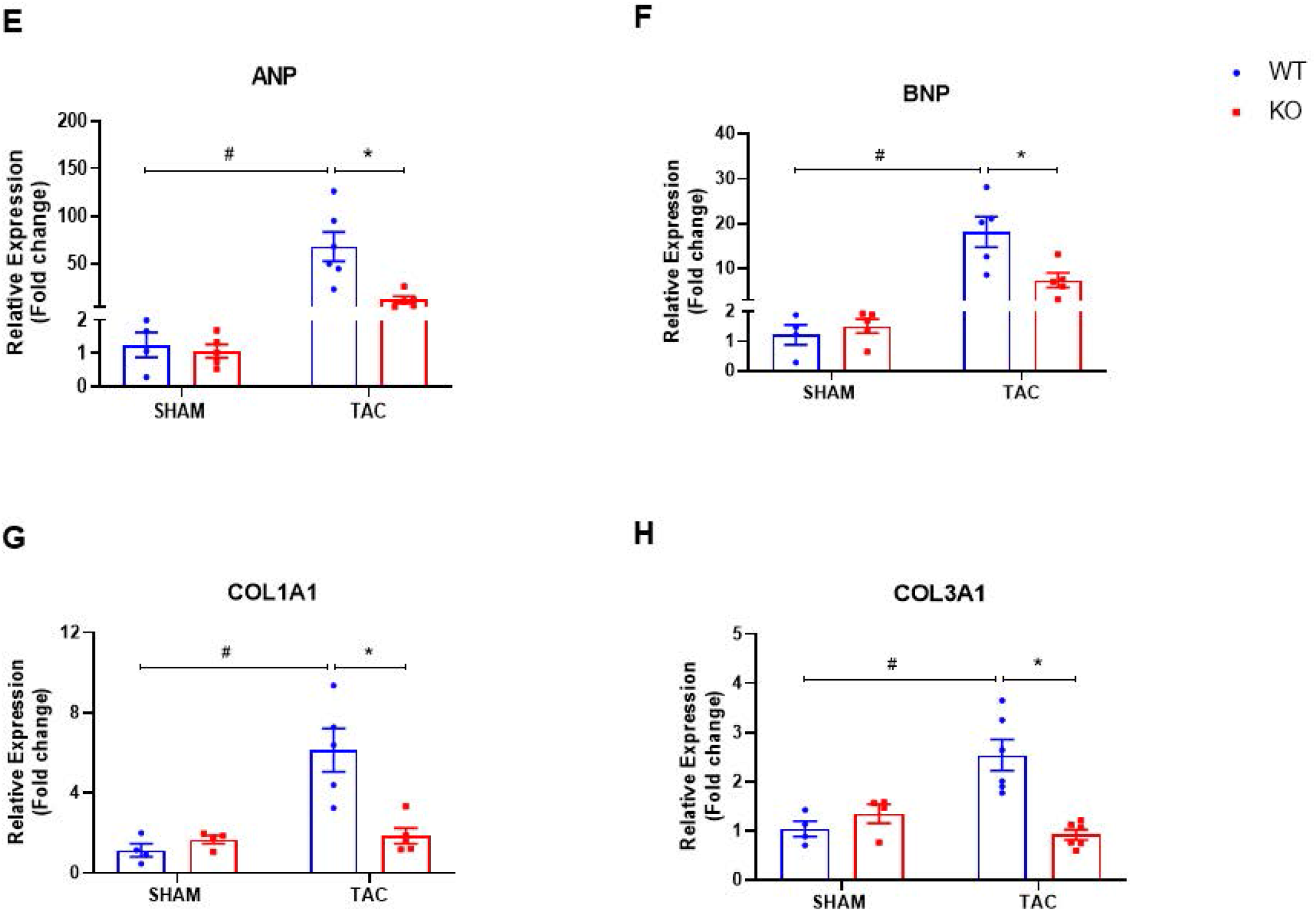
Deletion of CF-GSK-3α from resident cardiac fibroblasts prevents pressure overload-induced adverse cardiac remodeling. Morphometric studies were performed at 8 weeks after TAC surgery. Assessment of cardiac hypertrophy; **(A)** Heart weight (HW) to tibia length (TL) ratio and **(B)** Quantification of cardiomyocyte cross-sectional area (CSA). Assessment of cardiac fibrosis by Masson’s Trichrome staining; **(C)** Representative Trichrome-stained LV regions and **(D)** Quantification of LV fibrosis. Scale bar = 30 μm. RNA was extracted from the left ventricle of experimental animals and gene expression analysis was carried out by qPCR method; **(E)** ANP, **(F)** BNP, **(G)** COL1A1, and **(H)** COL3A1. Data were analyzed using one-way ANOVA followed by Tukey’s *post hoc* analysis and represented as mean ± SEM. N=4-9 per group. ^#^*P*<0.05 for WT-SHAM vs WT-TAC, \**P*<0.05 for WT-TAC vs.KO-TAC.

### Deletion of GSK-3α in myofibroblasts delays injury-induced cardiac remodeling & dysfunction

Cardiac injury triggers activation of fibroblast by the process called myofibroblast transformation. Myofibroblast plays a key role in the healing phase of injury. However, persistent survival of these cells even after injury resolution leads to pathological fibrotic remodeling and cardiac dysfunction. To define the role of GSK-3α in myofibroblast function we deleted GSK-3α specifically in myofibroblast using tamoxifen (TAM)-inducible periostin promoter-driven MerCreMer transgene (Postn-MCM). We crossed Postn-MCM mice with GSK-3α ^fl/fl^ mice to obtain GSK-3α ^fl/fl^ Postn^MCM^ mice. GSK-3α^fl/fl^/MCM^+/-^/TAM mice were conditional knockout (KO), whereas littermates GSK-3α ^fl/fl^/MCM^-/-^/TAM represented wild type controls (WT). The tamoxifen (TAM) chow diet protocol was employed to induce the FB-specific expression of Cre recombinase, as previously reported.^8^ To confirm the CF-specific deletion of GSK-3α, fibroblasts were isolated from the heart of experimental animals and proteins were extracted. Western blot analysis confirmed that TAM treatment led to a significant reduction of GSK-3α protein in the CFs from KO mice compared with littermate controls **(Figure 3A–3B)**.

**Figure 3:**
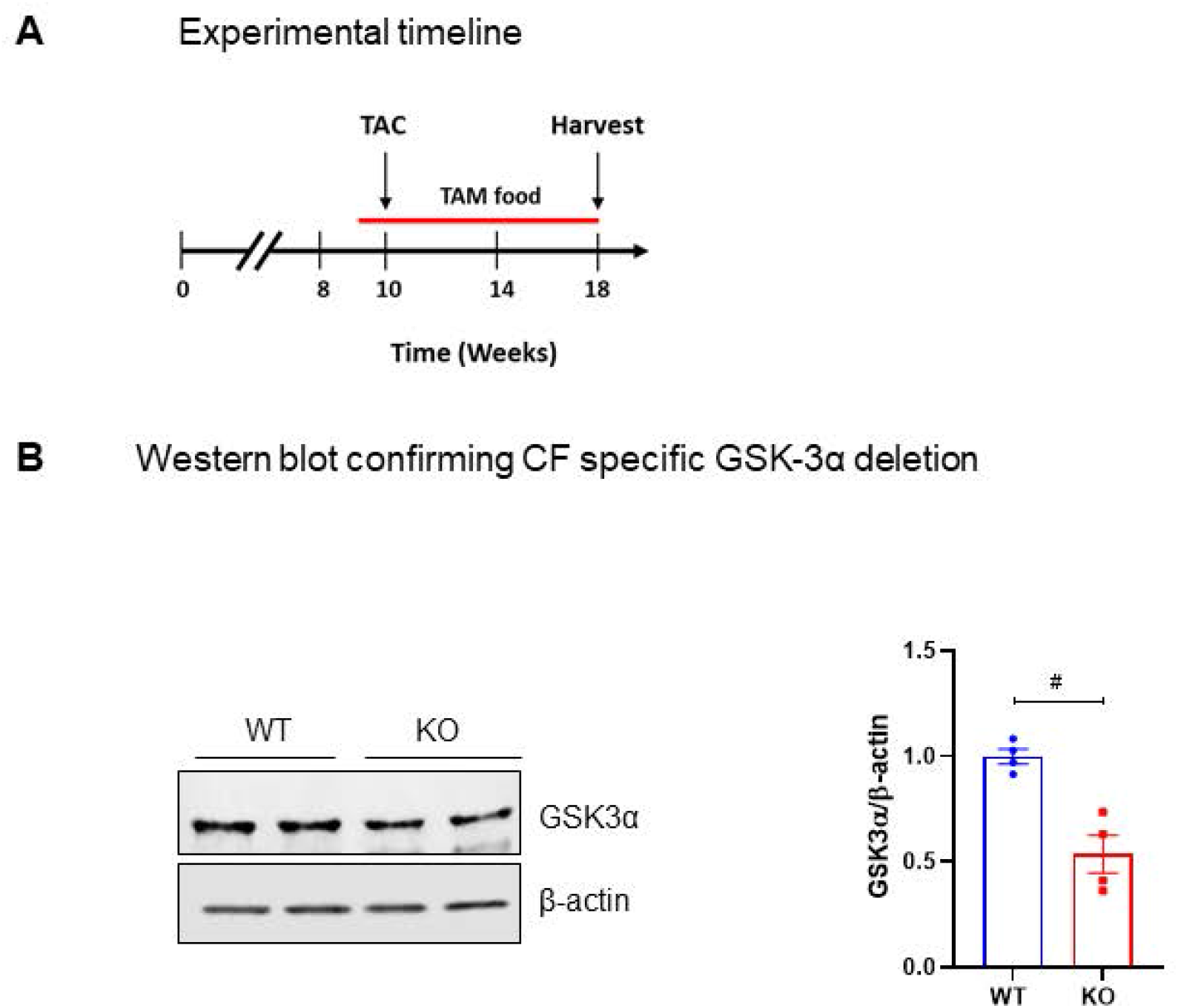
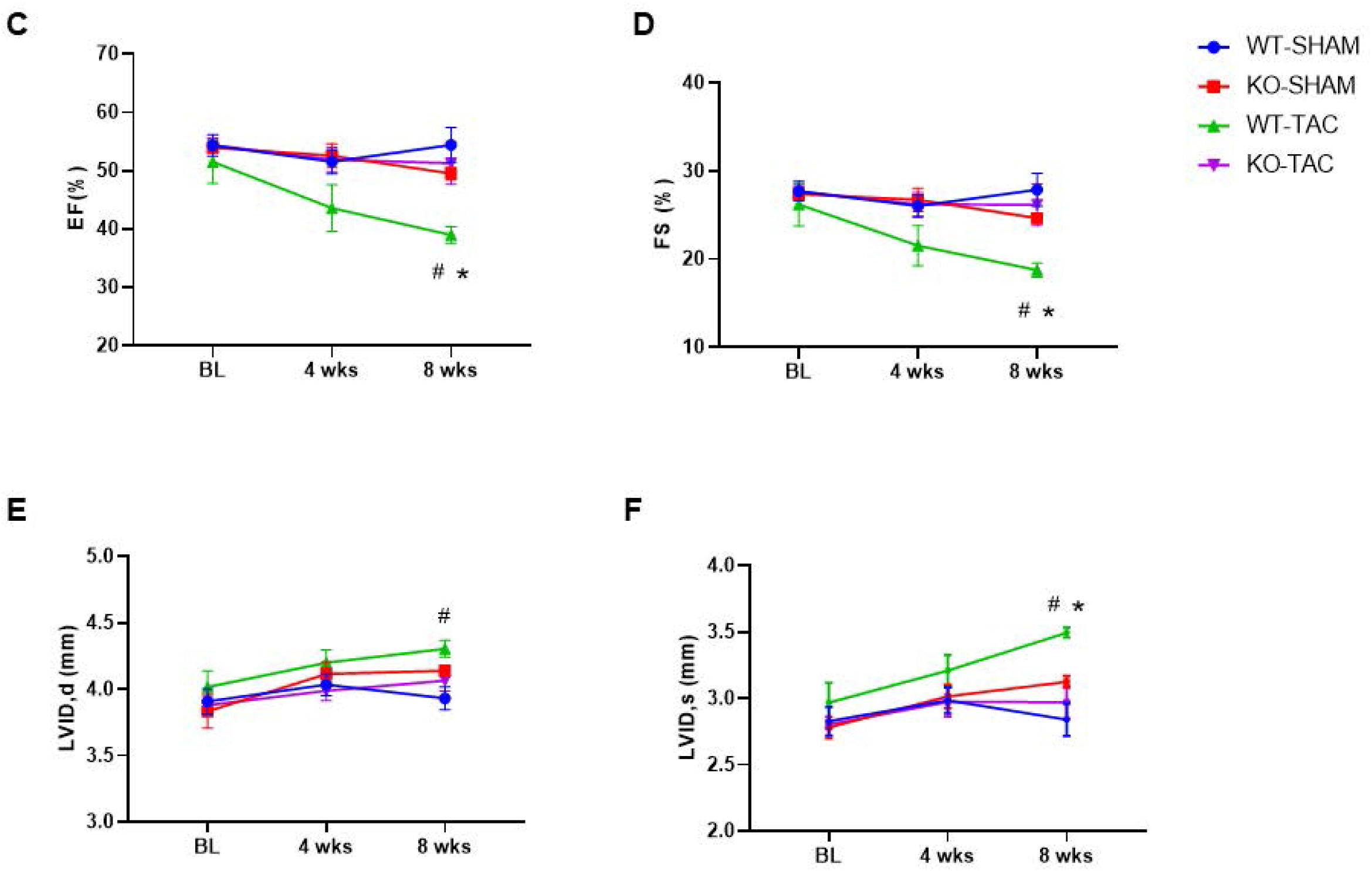
Deletion of CF-GSK-3α from myofibroblasts prevents pressure overload-induced cardiac dysfunction. **(A)** Experimental design. Two months old mice were fed the tamoxifen (TAM) chow diet. After 1 week of TAM treatment, mice were subjected to TAC surgery and are maintained on the TAM diet till the end of the study. **(B)** Western blot analysis of GSK-3α protein levels in cardiac fibroblasts at 4 weeks after TAM treatment. Data were analyzed using the nonparametric Mann-Whitney test and represented as mean ± SEM. N=4 per group. \**P*<0.05. Evaluation of cardiac function by m-mode echocardiography; **(C)** Ejection fraction (EF), **(D)** Fractional shortening (FS), **(E)** LV end diastolic interior dimension (LVID;d) and **(F)** LV end-systolic interior dimension (LVID;s). Data were analyzed using one-way ANOVA followed by Tukey’s *post hoc* analysis and represented as mean ± SEM. N=5-7 per group. ^#^*P*<0.05 for WT-SHAM vs WT-TAC, \**P*<0.05 for WT-TAC vs.KO-TAC.

After 1 week of TAM treatment, mice were subjected to trans-aortic constriction (TAC) surgery. We assessed cardiac function by serial echocardiography. The WT-TAC group showed lower LVEF and LVFS as compared to WT-SHAM confirming the development of cardiac dysfunction. Moreover, LVIDs were significantly increased in WT-TAC groups indicating the development of adverse cardiac remodeling. All these injury-induced adverse effects were remarkably prevented by myofibroblast specific GSK-3α deletion **(Figure 3C–3F)**.

To further verify the cardioprotective effects of myofibroblast specific GSK-3α deletion on cardiac remodeling, we harvested the heart of experimental animals at 8 weeks post-TAC for morphometrics and histological studies. The WT-TAC group showed a significant increase in HW/TL and CSA compared with the WT-SHAM group indicative of cardiac hypertrophy. These parameters were relatively lowered after myofibroblast-specific GSK-3α deletion **(Figure 4A & 4B)**. Masson’s Trichrome staining revealed excessive collagen deposition in WT-TAC groups and deletion GSK-3α from myofibroblast showed a major impact on cardiac fibrosis by normalizing it to that of the WT-SHAM group **(Figure 4C & 4D)**. Next, we analyzed the expression of signature genes related to cardiac hypertrophy (ANP and BNP) and cardiac fibrosis (COL1A1, COL3A1) using the qPCR method. As expected there was a significant increase in levels of these molecular markers in the WT-TAC group and myofibroblast specific GSK-3α deletion could normalize their levels **(Figure 4E–4H).** Taken together these findings suggest that deletion of GSK-3α post-injury affects myofibroblast function and delays injury-induced pathological remodeling.

**Figure 4:**
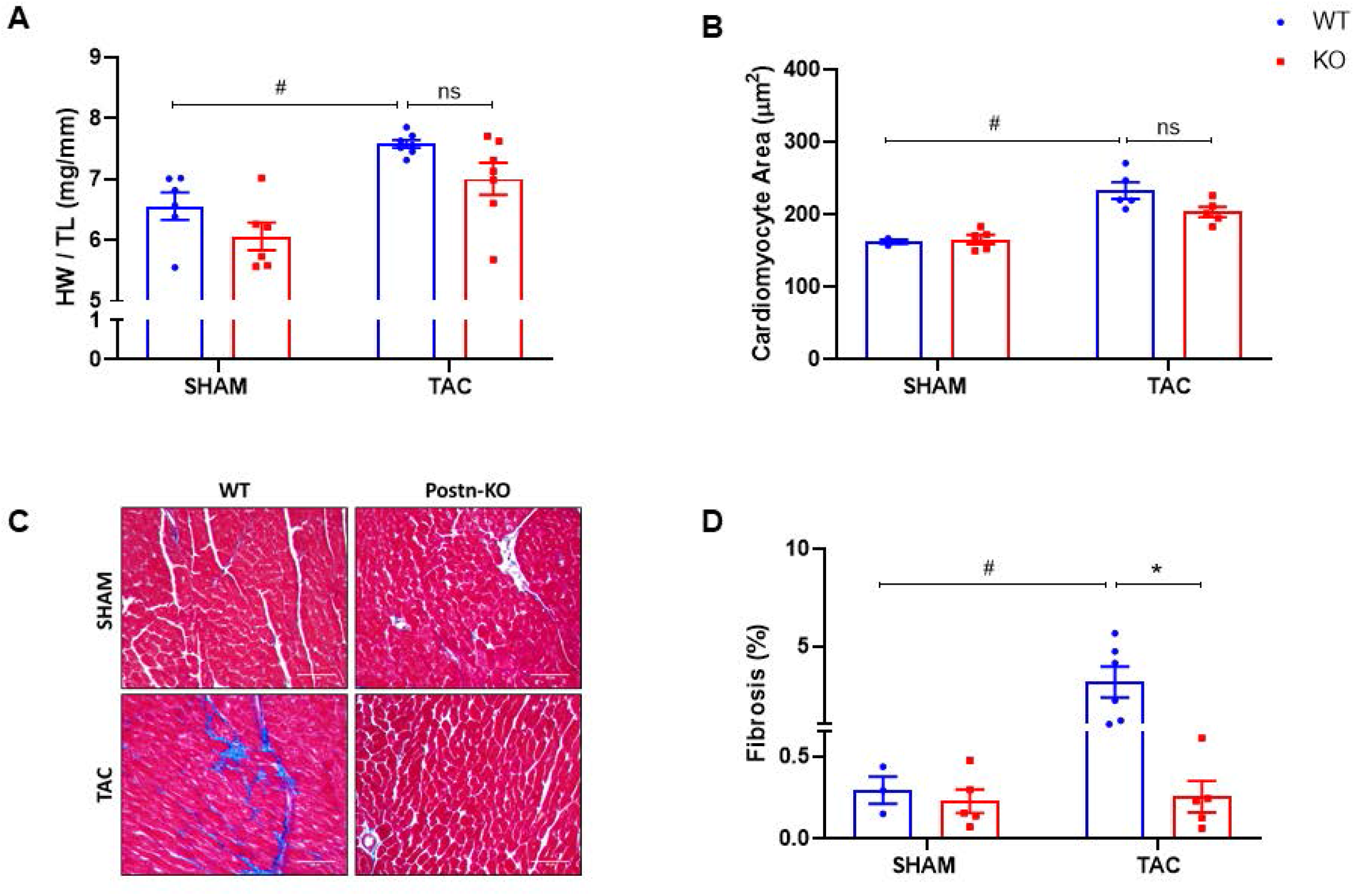
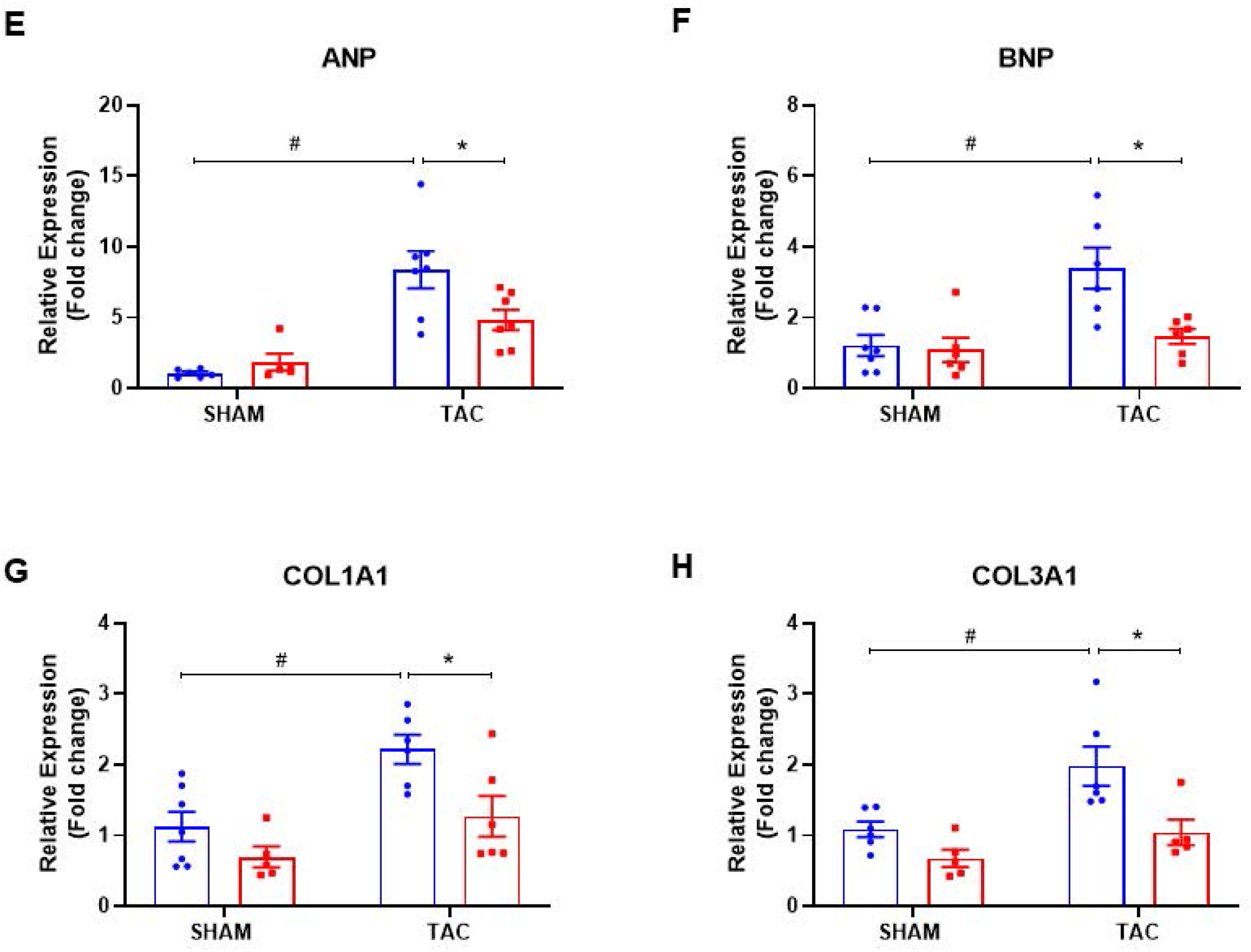
Deletion of CF-GSK-3α from myofibroblasts prevents pressure overload-induced adverse cardiac remodeling. Morphometric studies were performed at 8 weeks after TAC surgery. Assessment of cardiac hypertrophy; **(A)** Heart weight (HW) to tibia length (TL) ratio and **(B)** Quantification of cardiomyocyte cross-sectional area (CSA). Assessment of cardiac fibrosis by Masson’s Trichrome staining; **(C)** Representative Trichrome-stained LV regions and **(D)** Quantification of LV fibrosis. Scale bar = 30 μm. RNA was extracted from the left ventricle of experimental animals and gene expression analysis was carried out by qPCR method **(E)** ANP; **(F)** BNP; **(G)** COL1A1, and **(H)** COL3A1. Data were analyzed using one-way ANOVA followed by Tukey’s *post hoc* analysis and represented as mean ± SEM. N=5-7 per group. ^#^*P*<0.05 for WT-SHAM vs WT-TAC, \**P*<0.05 for WT-TAC vs.KO-TAC.

### CF-GSK-3α mediates pro-fibrotic effects independent of SMAD3

To define the role of GSK-3α in fibrotic remodeling, we examined the effect of GSK-3α deletion on myofibroblast transformation, a key event in fibrosis. The myofibroblast exhibits hypercontractile nature and significantly express α-SMA. Thus, to validate the phenotype we did functional assays such as cell migration and gel contraction assays which are based on these key features. To induce myofibroblast transformation we treated WT and GSK-3α KO mouse embryonic fibroblasts (MEFs) with TGF-β1 (10 ng/mL, 48h). We observed a significant reduction in cell migration, collagen gel contraction, and α-SMA expression in TGF-β1 treated GSK-3α KO MEF confirming that GSK-3α is required for myofibroblast transformation **(Figure 5A–5F)**.

**Figure 5:**
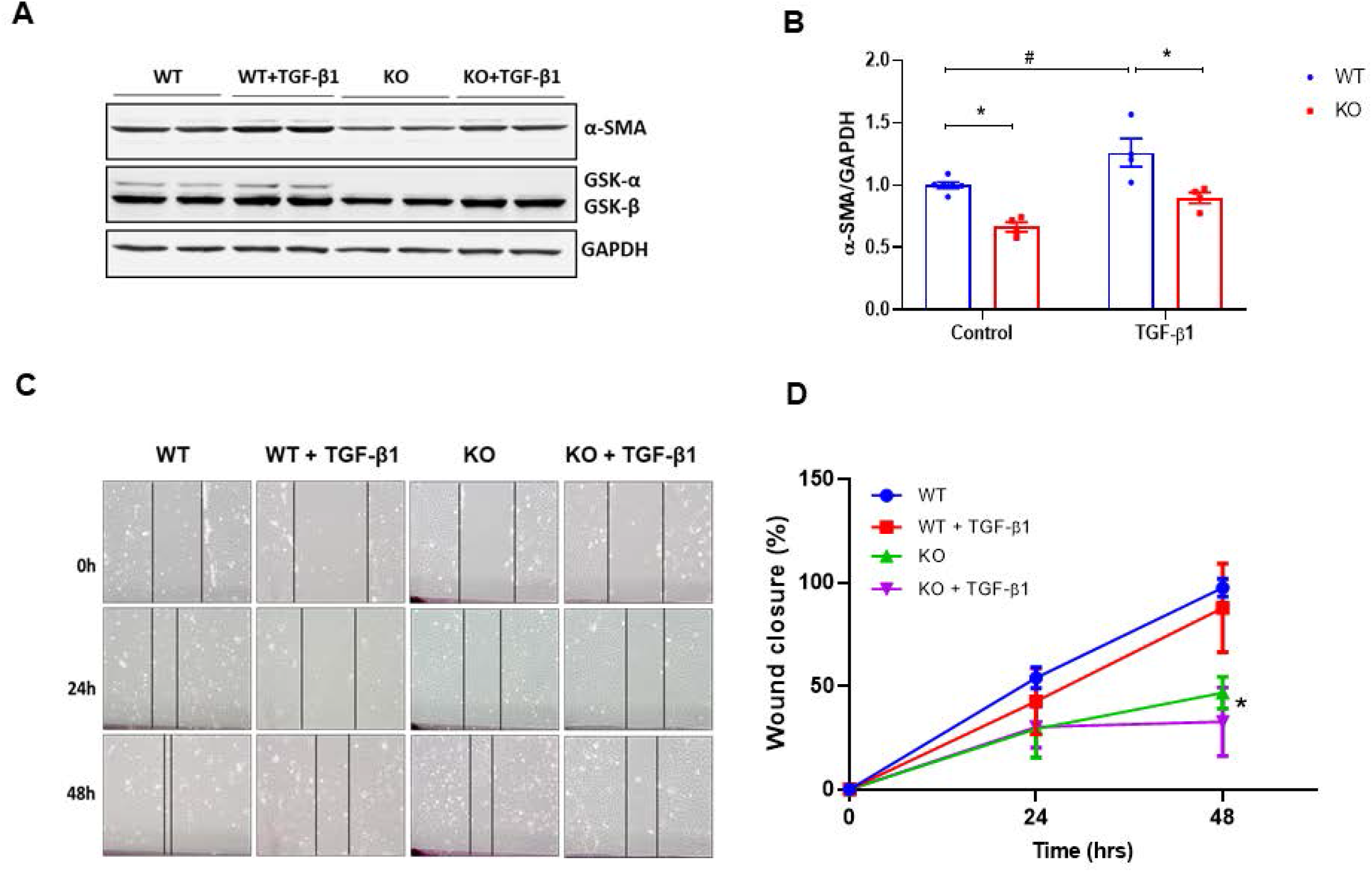
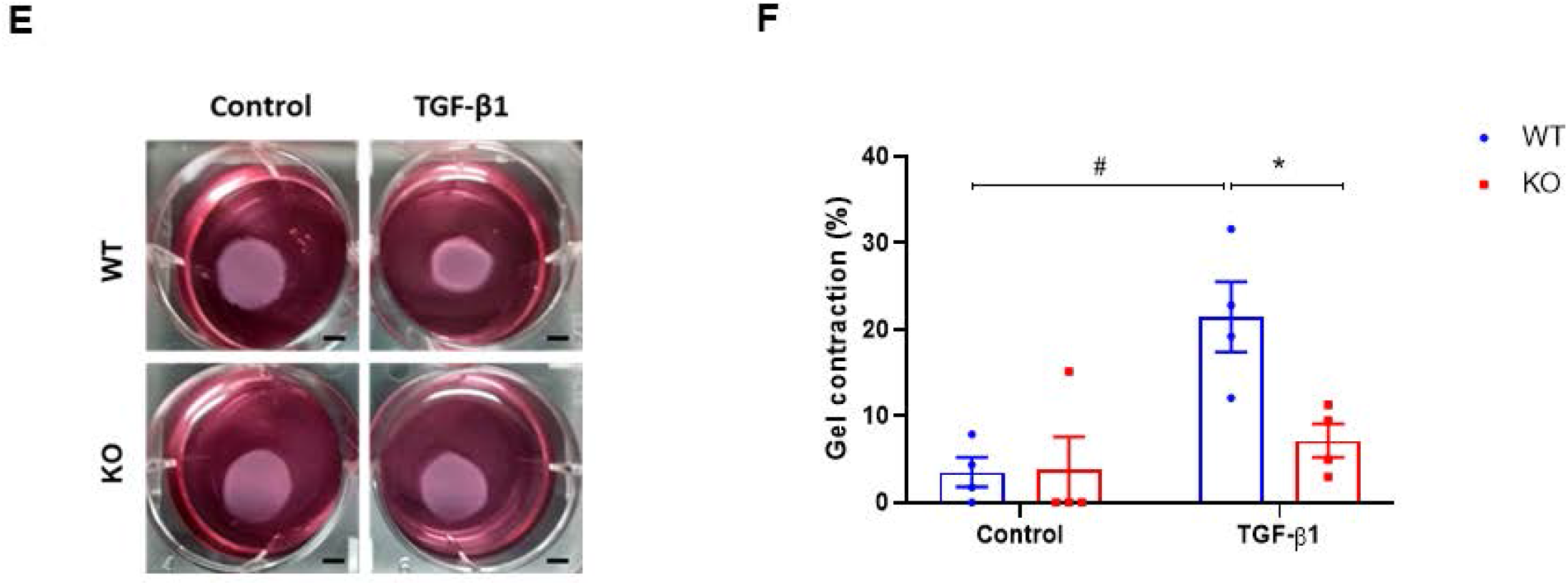
CF-GSK-3α regulates TGF-β1 induced myofibroblast transformation. WT and GSK-3α KO mouse embryonic fibroblasts (MEFs) were treated with TGF-β1 (10 ng/mL). **(A)** Western blot analysis of α-SMA protein levels after 48h of TGFβ1 treatment; Representative blot and **(B)** Quantification. WT and GSK-3α KO mouse embryonic fibroblasts (MEFs) were seeded and grown till confluency. Scratch was made followed by TGF-β1 (10 ng/mL) treatment. Wound closure was monitored; **(C)** Representative images and **(D)** Quantification of wound closure. Collagen gels were populated with WT and GSK-3α KO mouse embryonic fibroblasts (MEFs) followed by TGF-β1 (10 ng/mL) treatment. % gel contraction was calculated at 48h; **(E)** Representative image and **(F)** Quantification of gel contraction. Data were analyzed using one-way ANOVA followed by Tukey’s *post hoc* analysis and represented as mean ± SEM. N=3 per group. ^#^*P*<0.05 for WT-Control vs WT-TGFβ1, \**P*<0.05 for WT-TGFβ1 vs.KO-TGFβ1.

We have reported a critical role of CF-GSK-3β in TGF-β1-SMAD3 signaling, a well-known pro-fibrotic pathway.^15^ To test whether GSK-3α modulates TGF-β1-SMAD3 signaling, we examined the effect of GSK-3α deletion on SMAD3 activation. WT and GSK-3α MEFs were treated with TGF-β1 (10ng/ml, 1h), and SMAD3 activation was confirmed by analyzing levels of SMAD3 phosphorylation and nuclear translocation. Surprisingly, TGF-β1 induced SMAD3 activation was comparable in both WT and GSK-3α KO MEF **(Figure 6A–6D)**. To further confirm this observation, we isolated cardiac FBs from adult GSK-3α ^fl/fl^ mice and adenovirally transduced with either LacZ (Ad-LacZ) as control or Cre recombinase (Ad-Cre) to delete GSK-3α. Western blot studies confirmed that Ad-Cre transduction led to an 80% reduction in GSK-3α protein after 48h. These cells were further treated with TGF-β1 (10ng/ml, 1h). In line with MEF studies, we did not observe a significant effect of GSK-3α deletion on TGFβ1 induced SMAD3 phosphorylation **(Figure 6E–6G)**. Next, we employed a gain of function approach in which we used neonatal rat ventricular fibroblasts (NRVF) isolated from 1- to 3-day-old rat pups and overexpressed GSK-3α adenovirally (Ad-GSK-3α). Consistent with loss of function studies, overexpression of GSK-3α did not affect the TGF–β1-induced SMAD3 activation **(Figure 6H–6J)**. Taken together, this data indicated the potential involvement of CF-GSK-3α in eliciting SMAD3 independent pro-fibrotic signaling.

**Figure 6:**
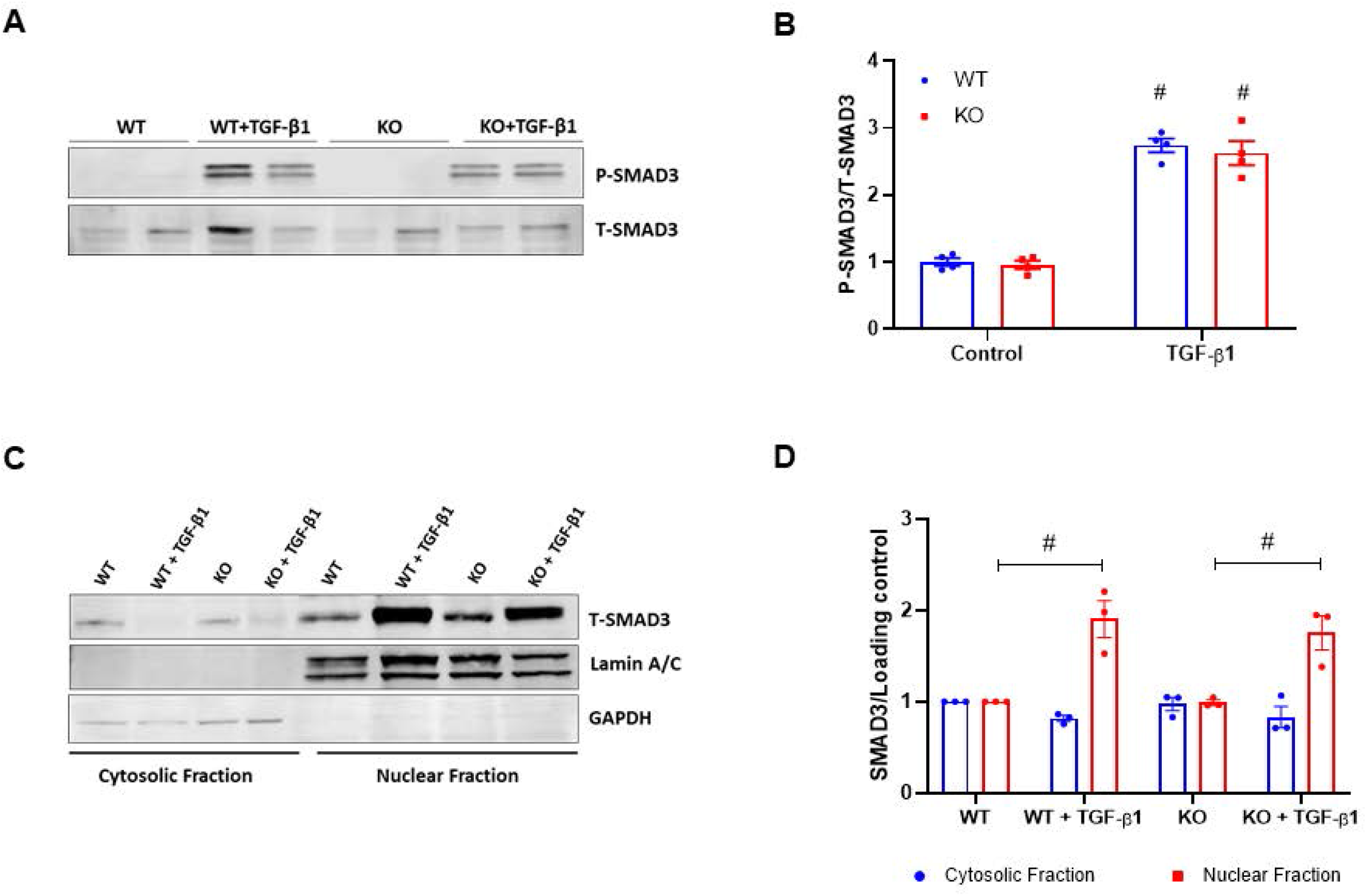
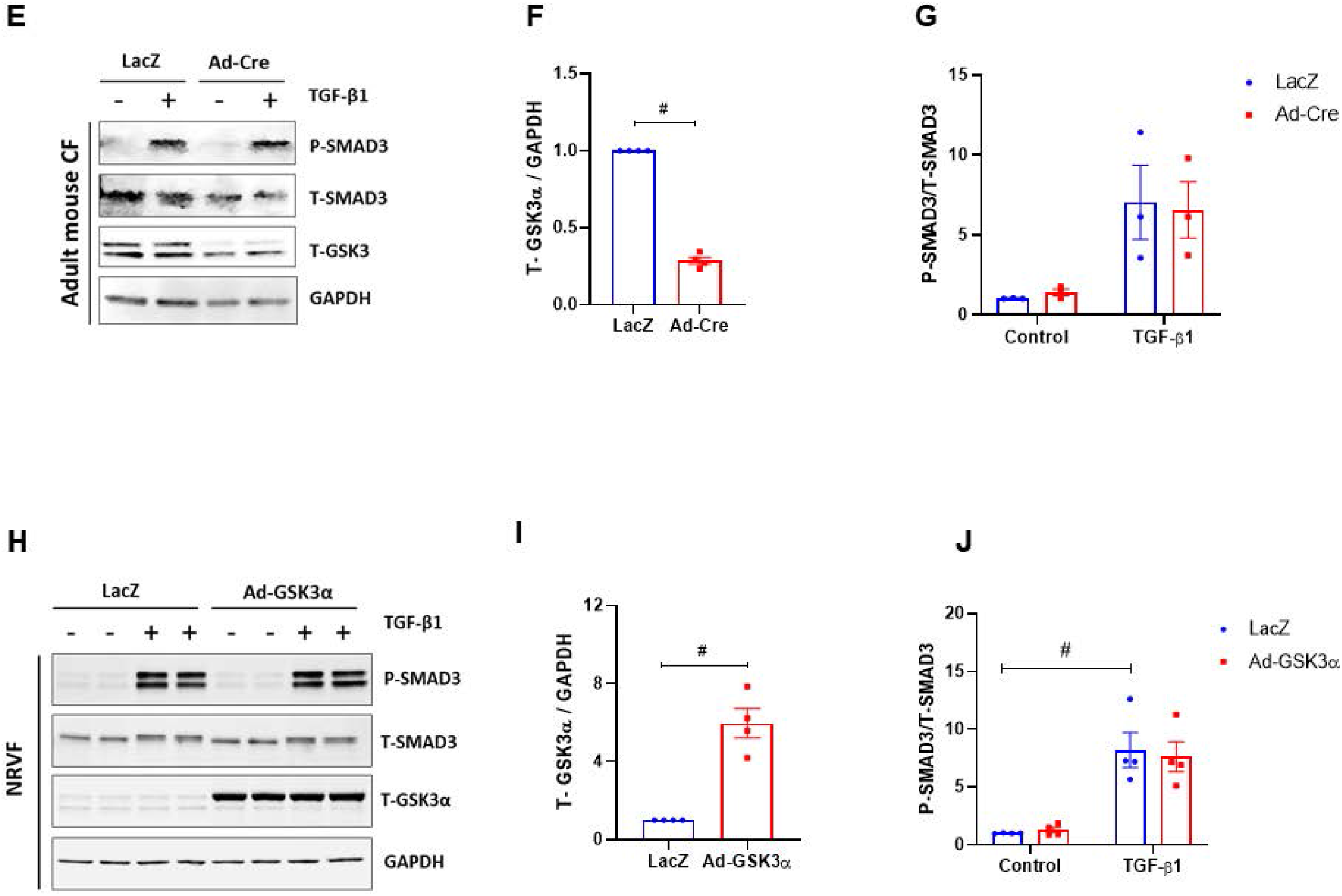
CF-GSK-3α mediates pro-fibrotic effects independent of SMAD3. WT and GSK-3α KO mouse embryonic fibroblasts (MEFs) were treated with TGF-β1 (10 ng/mL) for 1h. Western blot analysis of SMAD3 phosphorylation; **(A)** Representative blot and **(B)** quantification. Additionally, nuclear-cytoplasmic extraction was carried out and SMAD3 protein levels were analyzed by western blotting; **(C)** Representative blot, and **(D)** quantification. Adult CFs were isolated from GSK-3α ^fl/fl^ mice and GSK-3α was deleted by expressing Ad-Cre transfection. After transfection CFs were treated with TGF-β1 (10 ng/mL) for 1h. Western blot was carried out.; **(E)** Representative western blot and quantification of **(F)** GSK-3α and **(G)** SMAD3. For gain of function studies, GSK-3α was overexpressed in NRVFs, and cells were treated with TGF-β1 (10 ng/mL, 1h). Western blot was carried out.; **(H)** Representative western blot and Quantification **(I)** GSK-3α and **(J)** SMAD3. Data between the 2 groups were analyzed using the nonparametric Mann-Whitney test and represented as mean ± SEM. N=3 per group. ^#^*P*<0.05. Data between more than 2 groups were analyzed using one-way ANOVA followed by Tukey’s *post hoc* analysis and represented as mean ± SEM. N=3 per group. ^#^*P*<0.05 for WT-Control vs WT-TGFβ1.

### High Throughput Profiling of GSK-3α Regulated Fibroblast Kinome Reveals downregulation of RAF kinases in KOs

To gain further mechanic insights, we performed an unbiased kinome profiling of fibroblasts from CF-GSK-3α KO and littermate controls. At 4 weeks post-injury, CFs were isolated from WT and KO animals and total proteins were extracted. The kinome profiling was performed with the PamStation^®^12 high throughput microarray platform. Unsupervised hierarchical clustering of peptide signals was performed in BioNavigator and is displayed as a heatmap **(Figure. 7A)**. For the prediction of upstream kinases, peptide lists within the clusters were analyzed using the Kinexus Kinase Predictor. The upstream kinase analysis identified a significant reduction in phosphorylation of peptide targets of RAF family kinase, CDKs, RSKs, and AKT in GSK3α-KO-CFs **(Figure 7B–7C)**. Next, we uploaded the parent protein Uniprot ID for each peptide to GeneGo MetaCore program for network modeling. The mapping of significantly altered kinases against literature annotated interactions led to a generation of ERK-centric networks **(Figure 7D)**.

**Figure 7:**
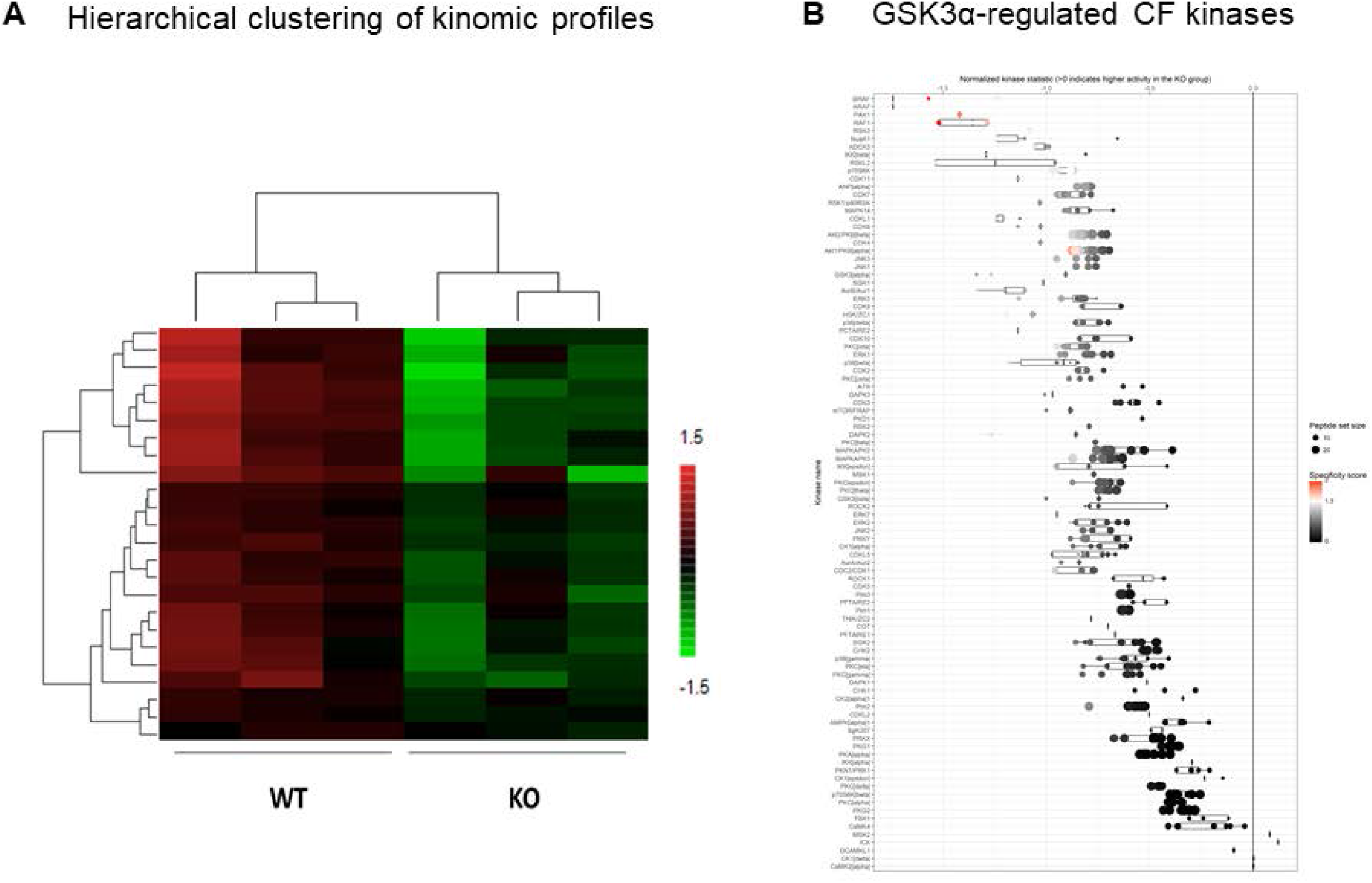
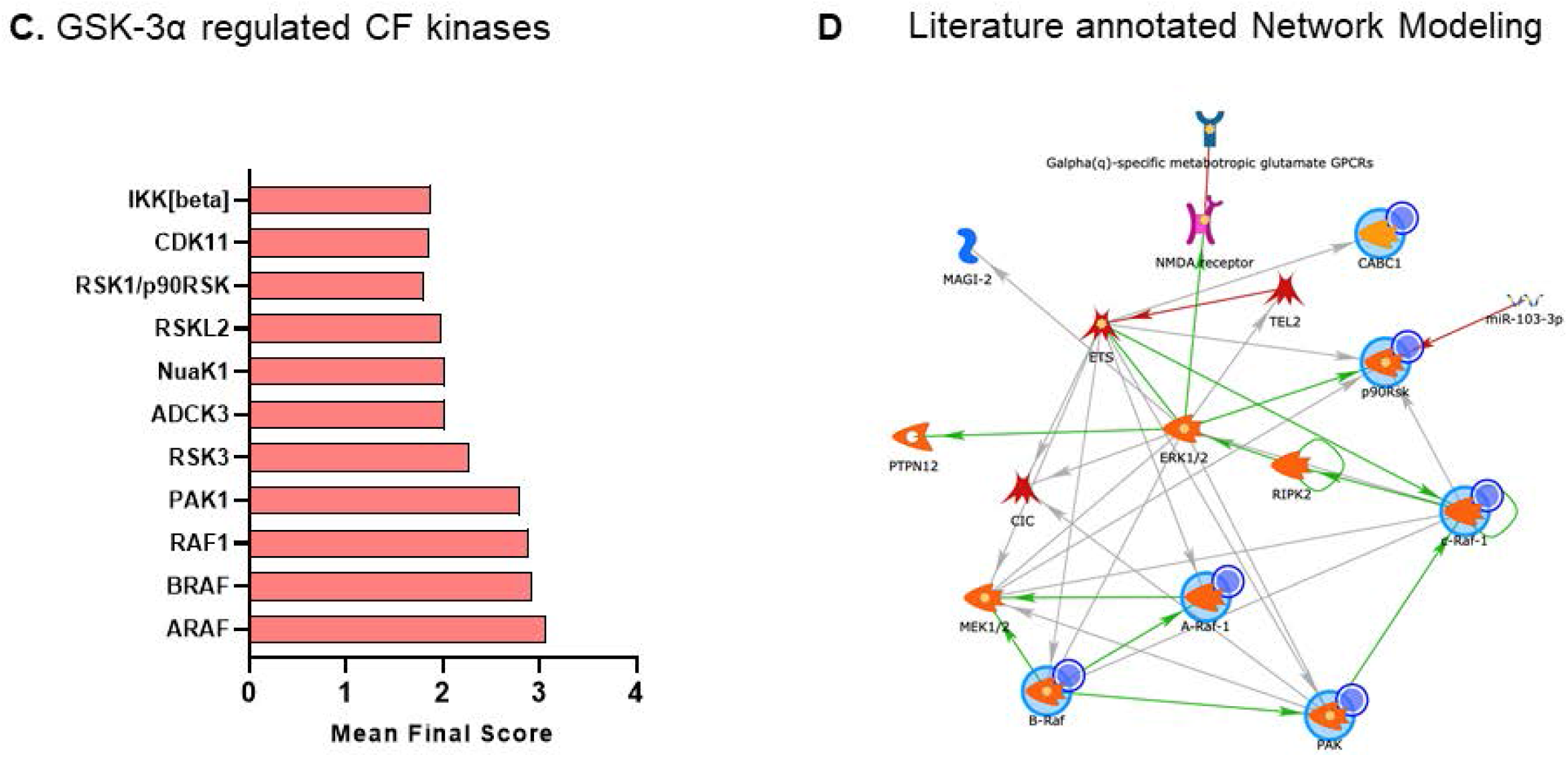
Profiling of GSK-3α regulated cardiac fibroblast Kinome. At 4 weeks post-TAC, proteins were extracted from CFs of WT and GSK-3α KO mice, and kinome profiling was carried out; **(A)** Hierarchical clustering of kinomic peptide phosphorylation signal intensity, **(B)** Bar plot displaying kinases identified using wholechip (all peptides) comparison for alteration in GSK-3α KO. The X-axis is the amount, and direction of kinases change, **(C)** Bar graph showing Mean Final Score of significantly altered kinases and **(D)** Literature annotated Network Modeling of significantly altered kinases. N=4.

### Effect of CF-GSK-3α deletion on IL-11 and ERK pathway

To validate our findings from kinome analysis, we isolated CFs from WT and KO mice at 8 weeks post-TAC and verified pERK levels by flow cytometry. As expected, we found a remarkable reduction in pERK^+^ CFs in KO mice **(Figure 8A)**. Next, to examine whether GSK-3α regulates the ERK pathway under pro-fibrotic conditions, we treated WT and GSK3α-KO cells with TGF-β1 (10 ng/mL, 10min) and assessed ERK activation by western blotting. Untreated GSK-3α KO MEF displayed considerably low levels of pERK as compared to WT. Additionally, TGF-β1-induced ERK phosphorylation was significantly inhibited in GSK3α-KO cells **(Figure 8B–8C)**.

**Figure 8:**
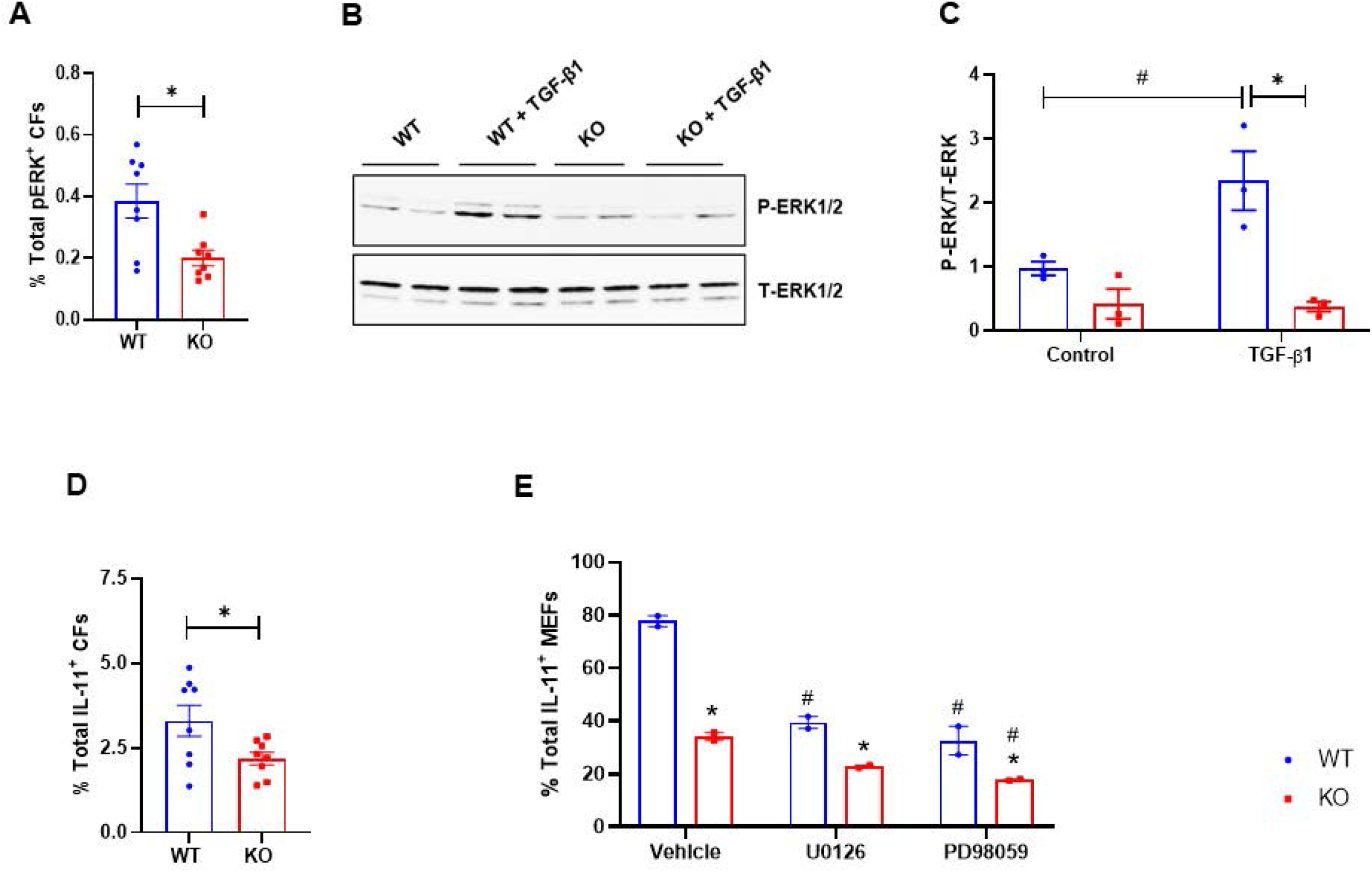
Effect of CF-GSK-3α deletion on IL-11 and ERK pathway. At 8 weeks post-TAC, CFs were isolated from WT and GSK-3α KO mice; **(A)** Flow cytometric analysis showing pERK +ve CFs (% total). Data between the 2 groups were analyzed using the nonparametric Mann-Whitney test and represented as mean ± SEM. N=3 per group. ^#^*P*<0.05. WT and GSK-3α KO MEFs were treated with TGF-β1 (10 ng/mL) for 10 min and ERK levels were assessed by western blotting; **(B)** Representative blot and **(C)** quantification. Data were analyzed using one-way ANOVA followed by Tukey’s *post hoc* analysis and represented as mean ± SEM. N=2 per group. ^#^*P*<0.05 vs WT-Vehicle, \**P*<0.05 WT vs KO. At 8 weeks post-TAC, CFs were isolated from WT and GSK-3α KO mice; **(D)** Flow cytometric analysis showing IL-11 +ve CFs (% total). Data between the 2 groups were analyzed using the nonparametric Mann-Whitney test and represented as mean ± SEM. N=3 per group. ^#^*P*<0.05. WT and GSK-3α KO MEFs were treated with ERK inhibitors (U0126, 10μM and PD98059, 5μM) for 24h **(E)** Flow cytometric analysis showing IL-11 +ve cells (% total). Data were analyzed using one-way ANOVA followed by Tukey’s *post hoc* analysis and represented as mean ± SEM. N=2 per group. ^#^*P*<0.05 vs WT-Vehicle, \**P*<0.05 WT vs KO.

Recent studies have identified IL-11 as a fibroblast-specific cytokine that mediates fibrogenic response downstream to diverse pro-fibrotic stimuli including TGF-β1. Importantly, IL-11 is demonstrated to act via an autocrine manner and is shown to cooperate with the ERK pathway to mediate fibrotic response in FBs. Hence, we examined the effect of GSK-3α deletion on IL-11 levels in CFs. Interestingly, our flow cytometric analysis results revealed a significant reduction in IL-11^+ve^ CFs in KO as compared to WT mice **(Figure 8D)**. To further validate the GSK-3α /IL-11/ERK crosstalk, we treated WT and KO GSK-3α MEF with ERK inhibitors and assess IL-11 levels. We observed low levels of IL-11 in vehicle-treated KO MEFs as compared to WT. ERK inhibitor treatment further reduced IL-11 levels in both cell types **(Figure 8E)**. These results indicate the potential role of GSK-3α /IL-11/ERK crosstalk in cardiac fibrosis under pathological conditions **(Figure 9)**.

**Figure 9:**
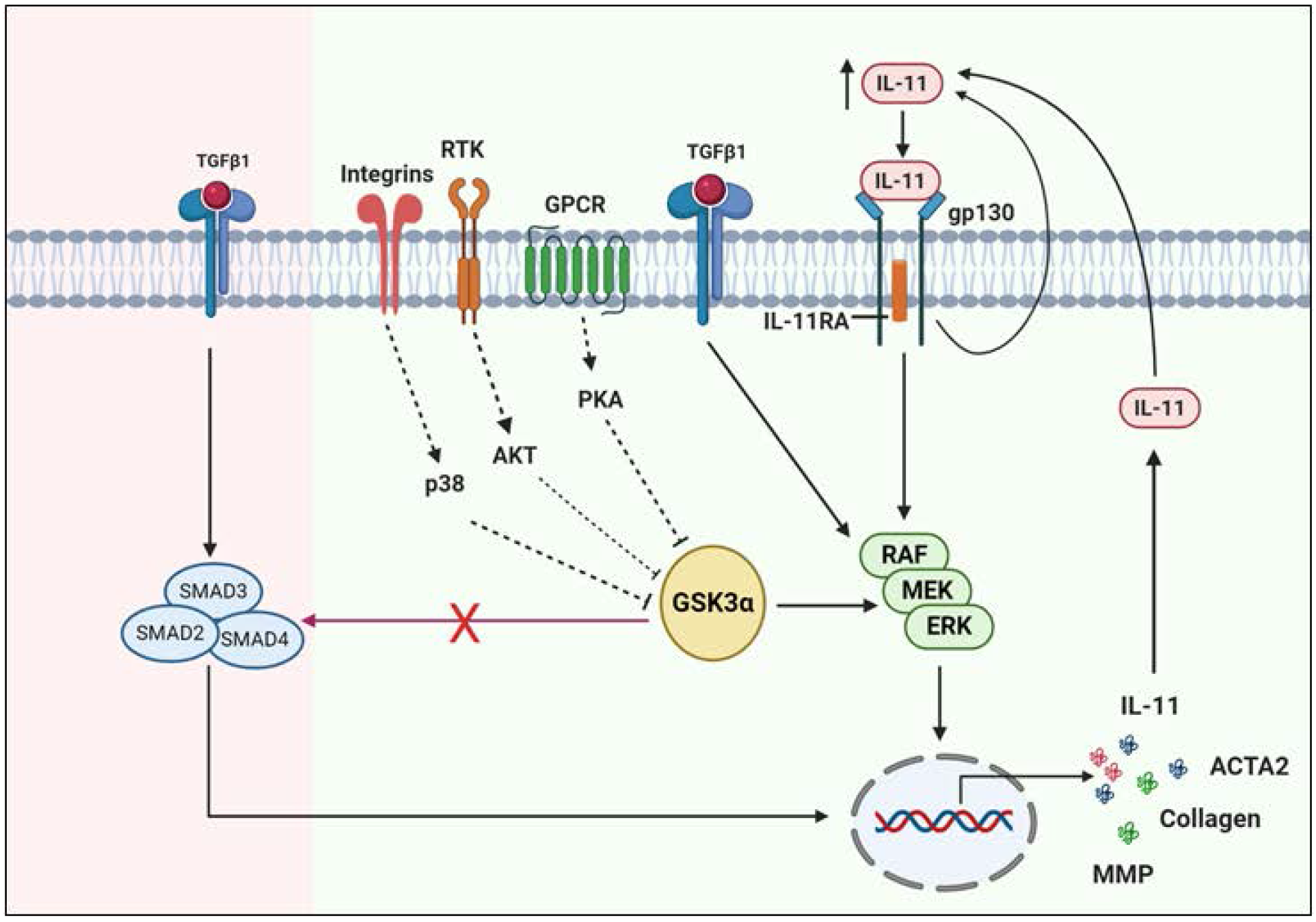
Schematic showing interactions of GSK-3α with IL-11 and ERK signaling pathways in cardiac fibroblast. CF-GSK-3α mediates profibrotic effects through IL-11 and ERK pathway in the injured heart while classical TGFβ1-SMAD3 signaling remained unaltered.

## Discussion

Herein, we discovered a critical role of CF-GSK-3α in the fibrotic remodeling of failing hearts. We employed two entirely novel inducible conditional FB-specific mouse models to demonstrate that CF-specific deletion of GSK-3α prevents injury-induced myofibroblast transformation, adverse fibrotic remodeling, and deterioration of cardiac function. We report that CF-GSK-3α mediates pro-fibrotic effects independent of the canonical TGFβ1/SMAD3 pathway. Furthermore, we demonstrate a causal role of CF-GSK-3α/IL-11/ERK crosstalk in regulating the fibrotic response to pathological stimuli.

In cardiac fibrosis, TGFβ1/SMAD3 signaling plays a crucial role.^9, 28^ Our lab has shown that in ischemic heart, GSK-3β deletion aggravates fibrosis via hyperactivation of SMAD3.^15^ Considering this observation, we were assuming that in the CF-GSK-3α KO heart, fibrosis might have reduced because of the direct effect of GSK-3α deletion on the TGFβ/SMAD3 pathway. To our complete surprise, GSK-3α KO CFs did not display alteration in TGFβ induced SMAD3 activation. This discrepancy could be explained by the well-known fact that GSK-3 isoforms play unique biological roles despite having great sequence homology.^17^ One of the best examples of the differential biological effect of GSK-3 isoforms is a global deletion of GSK3β is embryonic lethal while GSK-3α deletion does not affect embryonic development and mice survive several years.^20, 29^ Consistent with our findings, Guo et al. have demonstrated that GSK-3β, but not GSK-3α regulates the stability of non-activated SMAD3 and determines cellular sensitivity to TGFβ. This evidence suggests the isoform-specific substrate preferences could be one of the reasons for the differential effect of CF-GSK-3 isoforms on the TGFβ/SMAD3 pathway.

To identify the underlying mechanism of cardiac phenotype, we took an unbiased approach of GSK-3α regulated fibroblast kinome profiling. We found downregulation of RAF family kinases activity in GSK-3α KO CFs. Since the RAF-MEK-ERK pathway is implicated in fibrotic diseases across multiple organs.^31–35^ Hence we verified GSK-3α and ERK interaction and its relevance in the development of cardiac fibrosis. Thum et al. have shown that in pressure overloaded mouse heart microRNA-21 stimulates the ERK–MAP kinase signaling pathway in cardiac fibroblasts and contributes to disease pathogenesis.^36^ Moreover, recent studies from Ng et al. group have demonstrated that in response to a pro-fibrotic stimulus, IL-11 regulates fibrogenic protein synthesis via ERK signaling^16^ However, there are very few reports on GSK-3 and ERK interaction. An elegant study from Ding et al. group demonstrated a critical role of ERK in priming and inactivation of GSK-3β in cancer cells.^37^ In the case of cardiac pathophysiology, Sadoshima’s lab reported that in pressure overloaded hearts, cardiomyocyte (CM) specific GSK-3α overexpression leads to cardiac hypertrophy via inhibition of ERK, and when this inhibition is relieved by constitutively active MEK1 most of the pathological features can be rescued except cardiac fibrosis.^38, 39^ Our study suggests that CF-GSK-3α promotes fibrosis via IL-11 and ERK pathway in pressure overloaded hearts and highlights distinct fibroblast specific response of GSK-3α under similar pathological stimuli. Yet, the precise mechanism by which GSK-3α regulates IL-11 and ERK signaling in fibroblasts needs further studies.

The literature available on the role of GSK-3 isoforms in cardiac diseases suggests that the consequences of cardiac-specific GSK-3β manipulation are complex and context-dependent.^15, 18–21^ However, GSK-3α deletion has consistently shown cardioprotection irrespective of pathological stimuli.^40^ Our lab has reported that the CM-specific deletion of GSK-3α limits adverse remodeling and improves cardiac function post-MI.^23^ Furthermore, recently, we have demonstrated that the CM-specific deletion of GSK-3α protects from pressure overload-induced LV remodeling and cardiac dysfunction.^22^ Additionally, the critical contribution of GSK-3α to pathological cardiac hypertrophy is recapitulated in the gain of function studies by employing transgenic and KI mouse models. ^19, 39^ Consistent with all these studies, here we also observed the beneficial effects of fibroblast specific GSK-3α deletion against pressure overload-induced cardiac damages. Taken together, these critical pre-clinical data provide a proof of concept that therapeutic inhibition of GSK-3α could be a viable target for heart failure management.

In summary, we report that CF-specific deletion of GSK-3α prevents adverse fibrotic remodeling and improves the function of the diseased heart. Mechanistically, we have demonstrated for the first time that in response to pro-fibrotic stimuli, GSK-3α cooperate with the IL-11 and ERK pathway to mediate the fibrogenic response. Clinically, given the profound findings of the present study and literature discussed above, selectively targeting GSK-3α could be a novel and safe strategy to limit cardiac fibrosis and heart failure.

## NON-STANDARD ABBREVIATIONS AND ACRONYMS

CF: Cardiac Fibroblast
FB: Fibroblast
NRVF: Neonatal Rat Ventricular Fibroblast
CSA: Cross-Sectional Area
HW: Heart weight
LV: Left Ventricle
LW: Lung weight
MCM: Mer-Cre-Mer
TAM: Tamoxifen
TL: Tibia length

