## Supplemental Tables for "Cardiac fibroblast GSK-3α mediates adverse myocardial fibrosis via IL-11 and ERK pathway"

Supplemental Table 1: Antibodies and dilutions used for Western blot analysis

| No. | Antibody | Vendor | Catalog No. | Dilution |
|---|---|---|---|---|
| 1 | p-ERK1/2 (Thr202/Tyr204) | Cell Signaling Technology | 4370S | 1:1000 |
| 2 | ERK1/2 | Santa Cruz Biotechnology | sc-93 | 1:1000 |
| 3 | pSMAD3 | Abcam | ab52903 | 1:1000 |
| 4 | Total SMAD3 | Abcam | Ab40854 | 1:1000 |
| 5 | GSK-3α/β | Cell Signaling Technology | 5676 | 1:1000 |
| 6 | GSK-3α | Cell Signaling Technology | 4337 | 1:1000 |
| 7 | GAPDH | Fitzgerald | 10R-G109a | 1:10000 |
| 8 | β-actin | Santa Cruz Biotechnology | sc-4778 | 1:1000 |
| 9 | α- SMA | Sigma | A5228 | 1:5000 |
| 10 | IRDye 680LT Goat anti-Rabbit IgG | LI-COR Biosciences | 926-68021 | 1:3000 |
| 11 | IRDye® 800CW Goat anti-Mouse IgG | LI-COR Biosciences | 925-32210 | 1:3000 |

Supplemental Table 2: TaqMan gene expression assays and control used for qPCR analysis

| No. | Gene | Assay ID | Catalog No. | Vendor |
|---|---|---|---|---|
| 1 | COL1A1 | Mm00801666_g1 | 4331182 | Applied Biosystems |
| 2 | COL3A1 | Mm00802296_g1 | 4331182 | Applied Biosystems |
| 3 | ANP | Mm01255747_g1 | 4331182 | Applied Biosystems |
| 4 | BNP | Mm01255770_g1 | 4331182 | Applied Biosystems |
| 5 | Eukaryotic 18S rRNA Endogenous Control | | 4319413E | Applied Biosystems |

Supplemental Table 3: Antibodies used for flow cytometric analysis

| No. | Antibody | Vendor | Catalog No. |
|---|---|---|---|
| 1 | p-ERK1/2 (Thr202/Tyr204) | eBioscience | 53-9109-42 |
| 2 | MEFSK4 (feeder cell antibody) | Miltenyibiotec | 130-120-802 |
| 3 | CD31 | BD Pharmingen | 563356 |
| 4 | CD45 | BD Pharmingen | 557659 |
| 5 | Purified Rat Anti-Mouse CD16/CD32 | BD Pharmingen | 553141 |
| 6 | IL-11 | Santa Cruz Biotechnology | SC-133063 PE |
